# SPARClink: an interactive tool to visualize the impact of the SPARC program

**DOI:** 10.1101/2021.10.22.465507

**Authors:** Sanjay Soundarajan, Sachira Kuruppu, Ashutosh Singh, Jongchan Kim, Monalisa Achalla

## Abstract

The NIH SPARC program seeks to accelerate the development of therapeutic devices that modulate electrical activity in nerves to improve organ function. SPARC-funded researchers are generating rich datasets from neuromodulation research that are curated and shared according to FAIR (Findable, Accessible, Interoperable, and Reusable) guidelines and are accessible to the public on the SPARC data portal. Keeping track of the utilization of these datasets within the larger research community is a feature that will benefit data generating researchers in showcasing the impact of their SPARC outcomes. This will also allow the SPARC program to display the impact of the FAIR data curation and sharing practices that have been implemented. This manuscript provides the methods and outcomes of SPARClink, our web tool for visualizing the impact of SPARC, which won the 2nd prize at the 2021 SPARC FAIR Codeathon. With SPARClink, we built a system that automatically and continuously finds new published SPARC scientific outputs (datasets, publications, protocols) and the external resources referring to them. SPARC datasets and protocols are queried using publicly accessible REST APIs (provided by Pennsieve and Protocols.io) and stored in a publicly accessible database. Citation information for these resources is retrieved using the NIH reporter API and NCBI Entrez system. A novel knowledge-graph-based structure was created to visualize the results of these queries and showcase the impact that the FAIR data principles can have on the research landscape when they are adopted by a consortium.

## Introduction

The NIH Common Fund’s Stimulating Peripheral Activity to Relieve Conditions (SPARC) program aims to transform our understanding of nerve-organ interactions with the intent of advancing bioelectronic medicine towards treatments that change lives (1). The SPARC program employs a FAIR (Findable, Accessible, Interoperable, and Reusable) first approach for its datasets, protocols, and publications, hence enabling the data to be easily reused by the research communities globally. The SPARC Data Portal can be used as the gateway to access fully curated datasets at any time (2). Using the portal, researchers can search for data used in real-world experiments to verify or corroborate studies in device development. There is also potential for the data generated by the SPARC program to be useful outside the current field of study showcasing the benefits of multi-discipline data generation and sharing (3).

All SPARC datasets are curated by the researchers according to the SPARC Data Standards (SDS), a data and metadata structure derived from the Brain Imaging Data Structure (BIDS) (4). Several resources are made available to SPARC researchers for making their data FAIR, such as the cloud data platform Pennsieve, the curated vocabulary selector and annotation platform SciCrunch, the open-source computational modeling platform o^2^S^2^PARC, the online microscopy image viewer Biolucida, and the data curation software SODA (4–6). The datasets submitted by researchers also follow an extensive curation process where teams from the SPARC Data Resource Center (DRC) examine the submitted data and work with the researchers to ensure all aspects of the FAIR data principles are being followed (4,6,7). Once these datasets are made public, access to them is provided through the Pennsieve Discover service and sparc.science, the official access point of the SPARC Portal (8).

While the submission and curation of data are simplified with such tools, one of the larger benefits of the FAIR guidelines is the ability to reuse data in other studies by other researchers around the world. However, a researcher who has submitted a dataset might not always be aware of the reuse of their original submitted data since current citation indexing tools, like Google Scholar, do not account for datasets. To address this shortcoming, we developed SPARClink during the 2021 SPARC FAIR Codeathon (July 12th, 2021 – July 26th, 2021) (9), a system that queries all external publications using open source tools and platforms and creates a database and visualizations of citations that are helpful to showcase the impact of the SPARC consortium. In this instance, we define impact as the frequency of citations of SPARC-funded resources. By using citations as the key measure in SPARClink, we have created a method of showcasing the reuse of generated data and the benefits that FAIR data generation practices have on the overall scientific community. A visual representation of the reuse of data will allow both researchers and the general public to see the benefits of the concept of FAIR data and the immediate utilization of publicly funded datasets in advancing the field of bioelectronic medicine.

## Methods

Our solution can broadly be categorized into four steps. The first step involves the backend extraction of data using various APIs. The second step is setting up and storing the extracted data on a real-time database. The third step involves using machine learning to improve user experience by developing context-sensitive word clouds and smart keyword searches in the portal. The final step is used to create an engaging visualization that users of the SPARClink system will be able to interact with to view the extracted data. A visual representation of this workflow is shown in **Figure 1**.

**Figure 1.**
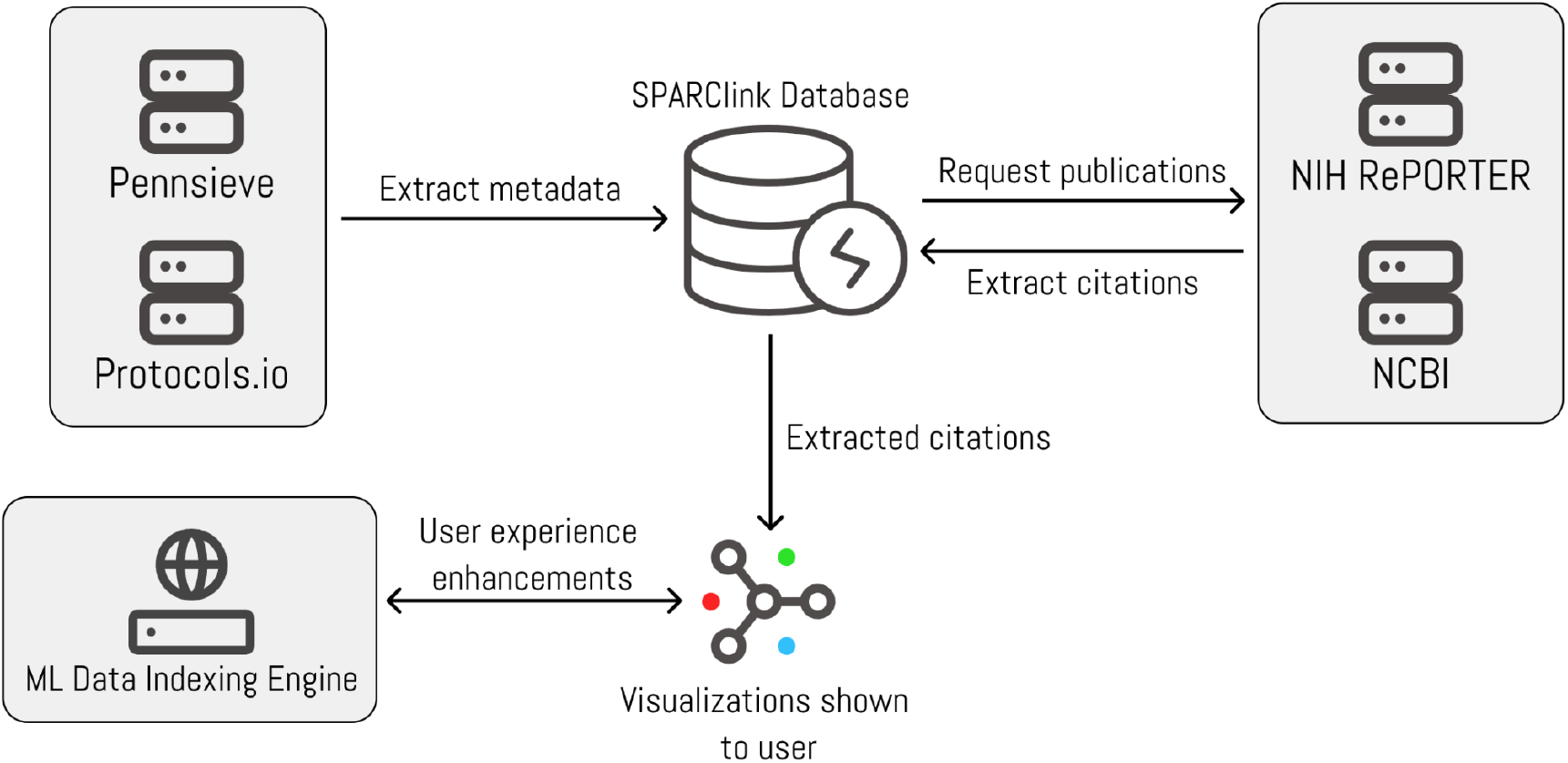
The flow of data between the submodules of SPARClink.

### 1. Extraction of data using APIs

We use the dataset information retrieved directly from the Pennsieve data storage platform by running the Pennsieve API to gather all publicly available SPARC datasets (10). The protocols stored on Protocols.io under the SPARC group are also queried via this method (11). A list of public and published DOIs is created in our database with additional information regarding the study authors and descriptions.

We use NIH RePORTER to retrieve data about the papers published as part of SPARC funding. Research articles that reference or mention these datasets, protocols, and publications are queried from NCBI (PubMed, PubMed central) repositories using the search endpoint of their Python API (12). **Figure 2** shows the overall flow of data between the APIs and resources queried to get the data. The NIH RePORTER API uses project number (also known as the award number) of NIH funding associated with SPARC datasets (this is provided by the author as additional metadata required when publishing a dataset) as an input to get details including a study identifier, name of the organization that received funding, country of the organization, amount of funding received and keywords of the project topic. The NCBI API uses an identifier for PubMed Central articles to retrieve information such as article name, journal name, year of publications, and authors.

**Figure 2.**
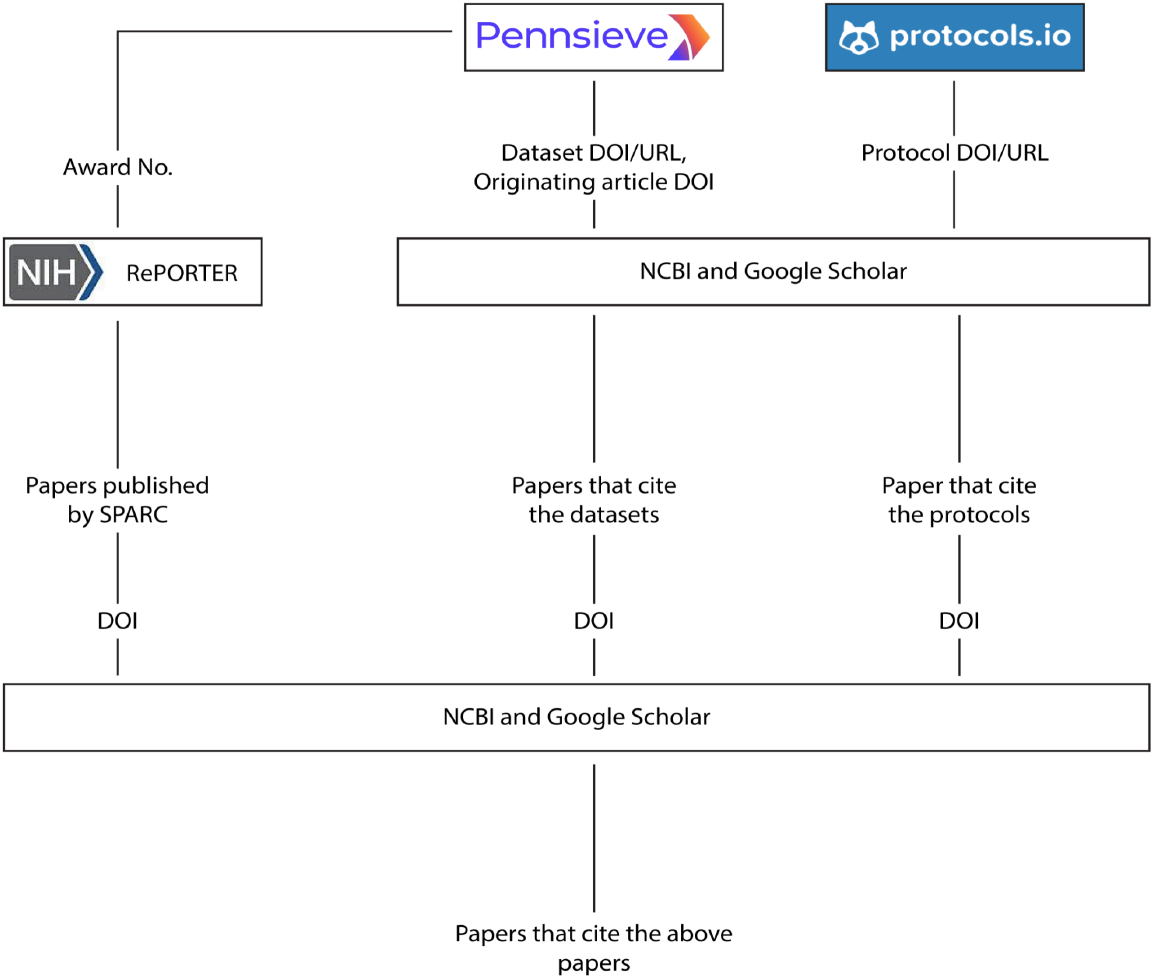
Methods implemented to gather citations of datasets, protocols, SPARC, and external publications

### 2. Storing extracted data in a database

We use Google’s Firebase real-time database to store all the information retrieved via the NIH RePORTER system. The data is stored in a JSON format with read access available to anyone via a dedicated URL. The data in this database is split up into four separate sections labeled Awards, Datasets, Publications, and Protocols. All the entries within this database are given a unique identifier. These identifiers are used to link the data within the database to form a relational database. The links within the data represent the citations or use of resources within other publications. All publications within the database are uniquely identified as either SPARC-funded publications or non-SPARC publications (external publications that cite SPARC datasets and publications.)

### 3. Displaying the extracted data to the user

The front-end demo of the SPARClink web page uses Vue.js to create a functional prototype of the SPARClink system. An interactive force-based undirected graph visualization was created using the D3.js javascript library. The choice of representing the results through such a graph was motivated by the desire to show an intuitively understandable way of showing the connected nature of citations and data reuse. The website itself is hosted on Vercel as a static front end (13). On the webpage, you can filter the visualizations by key terms or resource type to get a better understanding of the resources created using the SPARC program. A screenshot of the webpage is shown in **Figure 3**.

**Figure 3.**
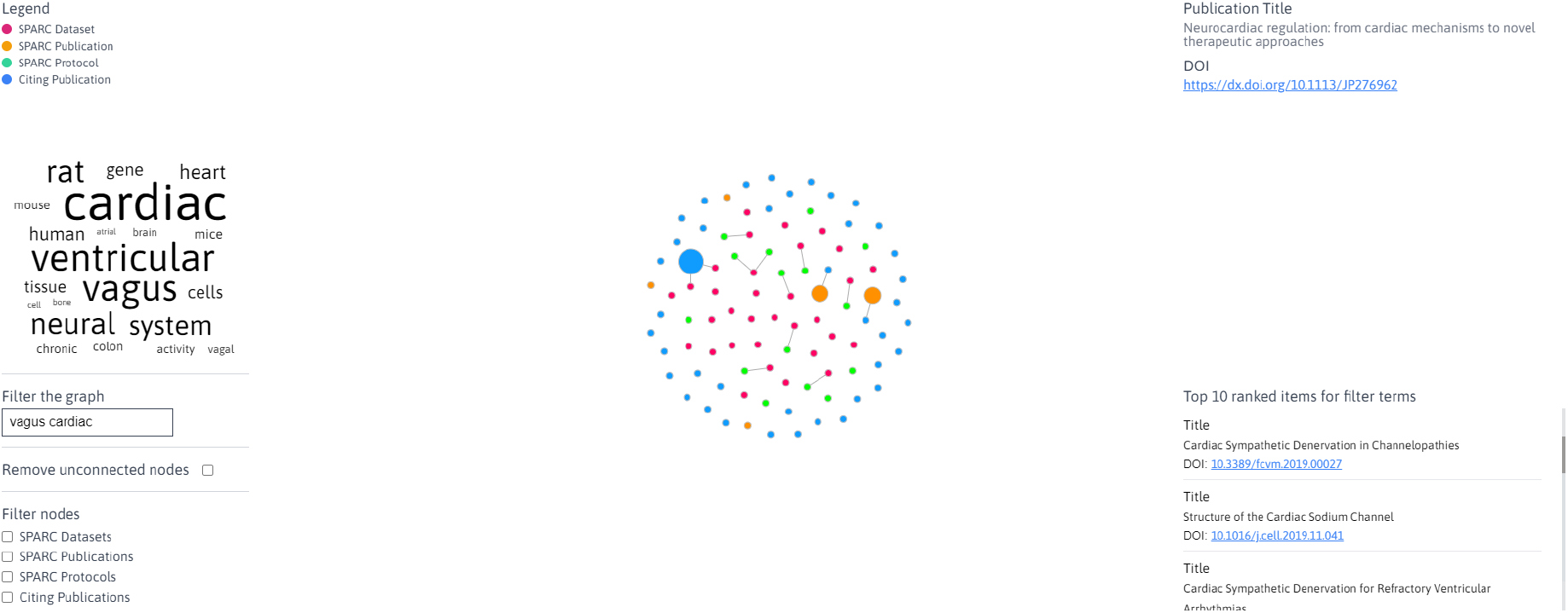
The design of the SPARClink webpage where the results from the machine learning module results are shown alongside the visualizations of SPARClink. The visualizations and the results in this figure have been filtered with the *vagus* and *cardiac* keywords.

### 4. Machine Learning Data Indexing Engine

To provide some additional functionality on the front-end demo of SPARClink we used machine learning algorithms to enhance the user experience. We call this side of the SPARClink project the Machine Learning Data Indexing Engine.

We use the Symspell algorithm present in the scikit learn package and trained it on the vocabulary built using the SPARClink database (14). We use delete-only edit candidate generation for generating different combinations of spelling errors and use both character level embedding and word embedding for recommending the most probable correct spelling. The output of the spell correction algorithm is used to generate sentence-level embedding and is then compared with the embeddings of different descriptions of the items in the dataset. We obtain a ranking of all the items in the dataset based on their similarity with the searched string. The top 10 are chosen to be shown on the front end.

This module is also used to generate keywords using the keyBERT pretrained model (15). It generates the top 50 keywords associated with the whole document. It also makes use of the Maximal marginal relevance algorithm to pick keywords that have a higher distance among them(16). This is to ensure diversity among the chosen keywords.

The engine also contains algorithms that learn vector embeddings of the descriptors of the elements present in the SPARClink database. Based on these vector embeddings the algorithms compute the similarity between the vector representation of each word in the vocabulary with the vector representing the whole dataset and find keywords that would describe the resource. A word cloud is generated based on the relevance of these results to further enhance the user experience.

## Results

Using SPARClink, researchers can aggregate all of the resources created through the SPARC program and quantify their impact. The visualization created by the SPARClink system is shown in **Figure 4**. The nodes in the undirected graph signify a unique SPARC resource (publication, protocol, and dataset) and the edges in the graph signify the citations or references as found by SPARClink. A well-connected graph of datasets and publications are observed but a significant number of protocols are seemingly distinct from the rest of the resources despite being pulled from the SPARC protocols.io group. This could be associated with protocols that are published on protocols.io but for which the associated datasets have not been made public yet.

**Figure 4.**
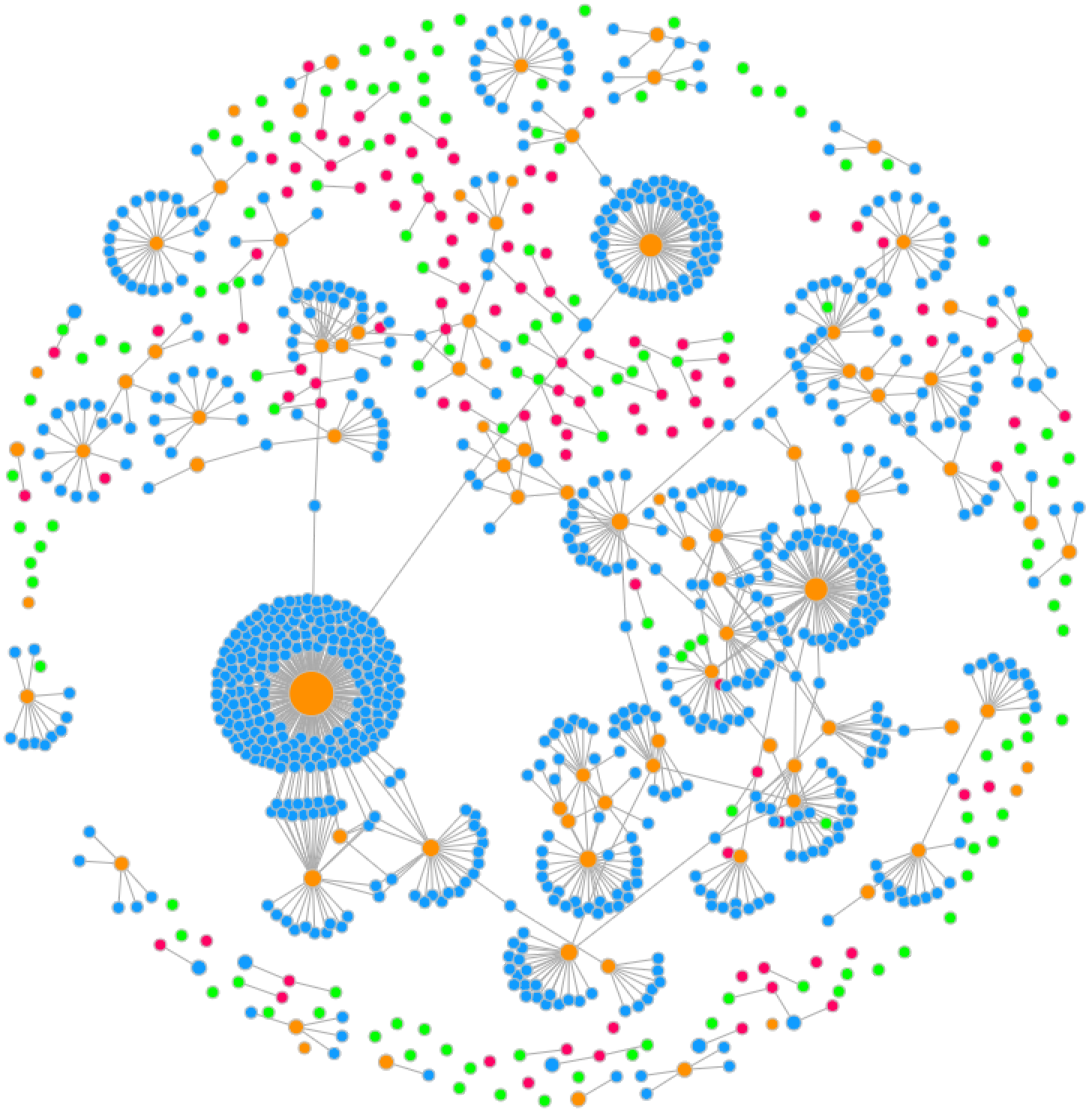
An interactive visualization created by SPARClink showing the connected nature of all of the SPARC resources.

The word map generated from the main dataset visualizations is shown in **Figure 5**. The size of the word with respect to its neighbors corresponds to the frequency and significance of the word within all the searchable metadata that we have indexed. Selecting any of the words in this map will automatically filter the SPARClink visualizations. Using a keyword filter on the graph will also prompt the top-ranking items for the keyword to be displayed on the side of the page. This ranking is shown as a scrollable list as shown in **Figure 6**. Both the word map and top-ranked recommendations are continuously updating themselves when new input terms are entered via the SPARClink webpage.

**Figure 5.**
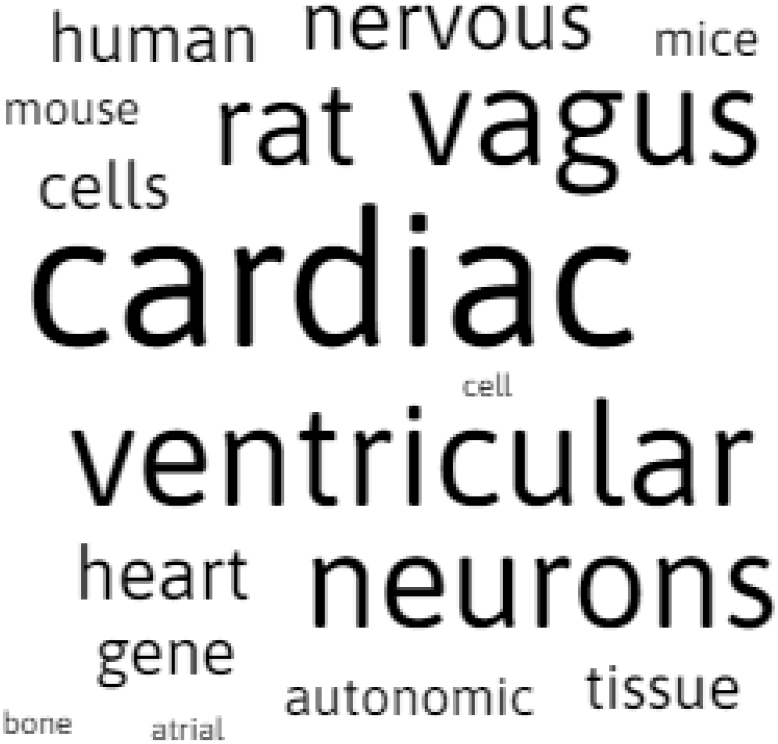
The word maps created by SPARClink are a visual representation of the most significant words shown in the graph-based visualization.

**Figure 6.**
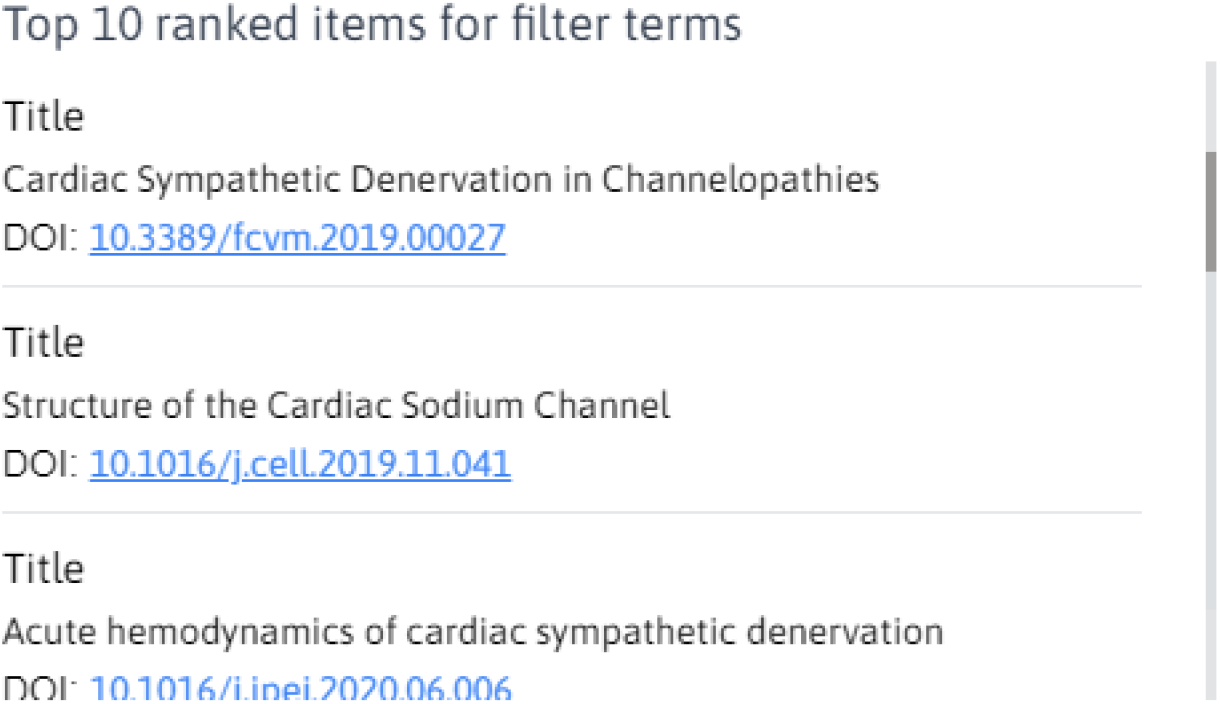
A list of resources that are recommended by SPARClink when a search term filter is provided by the user.

## Discussion and Conclusion

Using FAIR standards can greatly improve the use of data across multiple disciplines and potentially lead to new and exciting discoveries in the field of biomedical science. The benefits of employing the FAIR data principles for data generation, curation, and sharing can, however, be hard to quantify for researchers or members of the general public. Using a system like SPARClink, researchers at all levels can get up-to-date feedback on the use of their data and all the advantages that the FAIR standards provide in the area of advancing biomedical science. In this work, we developed such a tool for the SPARC program to enable quantification of the reuse of the FAIR SPARC resources (datasets, manuscripts, protocols).

The primary challenge in accomplishing this task lies in the fact that the SPARC datasets and protocols are not referenced in the bibliography of research manuscripts, which is the common practice. Instead, the SPARC dataset and protocol identifiers or URLs are only mentioned in the text or under supplementary materials, which makes querying this information a challenging task. Furthermore, datasets created in the SPARC program can be embargoed for up to 12 months to allow researchers enough time to document and publish their findings. However, protocols are made public immediately since protocols.io does not have an option to embargo the public publishing of these protocols. This could also add to the sparse graphs and we can expect the connectedness of this graph to improve as time goes on.

In the future, we plan on adding the Google Scholar system as an additional resource from where data is extracted. This should improve the connectedness of our extracted data network as well. Additional filtering functions and performance improvements for very large numbers of nodes are also planned. Currently, the tool is on an independent webpage but we also aim to integrate it directly within the SPARC portal such that visitors can conveniently visualize the reuse and impact of the different SPARC generated resources.

## Data and software availability

At the time of publication, the SPARClink system visualizations can be found at https://sparclink.vercel.app and are expected to be always online going forward. The backend system that queries all the publications is currently paused due to a lack of system resources. The code for SPARClink has been developed to be accessible to anyone who wants to fork the repository from GitHub and run a local version of this project. Instructions on how to run the modules locally are also available in the Github repository. The database of currently extracted citation data can be queried via REST protocols using the links provided below. The machine learning data indexing engine is hosted on a web server provided by pythonanywhere.com and is publicly accessible via its API endpoints. This module is also available to be run in local configuration seamlessly.

Source code available from: https://github.com/fairdataihub/SPARClink

Archived source code as at time of publication: https://doi.org/10.5281/zenodo.5550844

License: MIT

SPARClink extracted data can be queried at: https://sparclink-f151d-default-rtdb.firebaseio.com/

## Author Contributions

- This manuscript and the front-end webpage were prepared and created by Sanjay Soundarajan.
- Sachira Kuruppu developed the NCBI data extraction module.
- Ashutosh Singh worked on machine learning models and smart searching functions.
- Jongchan Kim developed the protocols.io data extraction module.
- Monalisa Achalla helped in the creation of this manuscript.

## Acknowledgments

We would like to thank the NIH Common Fund’s SPARC Program and the organizers of the 2021 SPARC FAIR Codeathon for their support during the development of this project.

